# Trained immunity in human monocyte enhances myeloid-T-cell pathogenic crosstalk in Ankylosing Spondylitis

**DOI:** 10.1101/2023.06.10.544264

**Authors:** Jinyi Zhao, Feng Liu, Claudia Worth, Davide Simone, Frank Penkava, Hui Shi, Jiaqi Li, Shristi Lama, Hai Fang, Jeong Seok Lee, Tae-Jong Kim, Lihua Duan, Paul Bowness, Liye Chen

## Abstract

Recent studies in infectious, cardiovascular and neurodegenerative diseases have established the presence of memory in innate immune cells. This “trained immunity (TI)” leads to an enhanced response to a second challenge. Monocytes in Ankylosing Spondylitis (AS), a common form of inflammatory arthritis, are known to be hyper-responsive to microbial stimulus lipopolysaccharide (LPS). We asked if TI is present in AS monocytes and, if so, how it contributes to disease pathology.

Using Single-cell RNA sequencing (scRNA-seq), flow cytometry and enzyme-linked immunosorbent assays (ELISA), we identify a subset of monocytes from AS patients exhibiting features of trained immunity and being hyperresponsive to LPS stimulation. Surprisingly, both trained monocytes in AS and β-glucan-trained monocytes from healthy donors are hyper-responsive to T-cell-induced activation. scRNA-seq of AS synovial mononuclear cells shows enrichment of a monocyte population with these/analogous features. Additionally, T cell-stimulated monocytes act back on T-cells to support Th17 responses (of established pathology in AS). Lastly, using genetic and chemical perturbations we show that ERN1, an AS risk gene enriched in this trained monocyte population, contributes to T-cell-induced monocyte activation.

Our data provide strong evidence for the first time for the key role of TI in common human inflammatory arthritis.

## INTRODUCTION

The term “trained immunity (TI)” was first proposed a decade ago for the concept that innate immune cells develop the memory of past insults (1). The presence of TI in myeloid cells previously exposed to microbial stimulus, vaccine or lipoprotein has now been well-established (2–4). Trained monocytes (and/or macrophages) exhibit epigenetic, transcriptional and metabolic changes leading to enhanced cytokine production upon a second innate stimulus (5–8). TI has been implicated in infectious, cardiovascular and neurodegenerative diseases but not inflammatory arthritis.

Ankylosing spondylitis (AS) is an immune-mediated inflammatory rheumatic disease with a strong genetic predisposition (9). The pathogenesis of AS is complex and still not yet fully understood, but a clear pathogenic role of interleukin (IL)-17A-producing T helper 17 (Th17) cells has been established. The frequency of Th17 cells is elevated in the peripheral blood of AS patients (10, 11), and biologics blocking IL-17A are effective treatments (12). Of note, IL-1β and IL-23 derived from myeloid cells are key upstream pro-inflammatory cytokines supporting the differentiation and expansion of Th17 cells (13–15). Interestingly, both IL-1β and IL-23 production are increased in myeloid cells from patients with AS (16, 17). Monocytes in AS patients show enhanced spontaneous (and stimulation-induced) production of IL-1β compared with healthy controls (16). In addition, both the level of serum IL-23 and the extent of lipopolysaccharides (LPS)-induced macrophage IL-23 production are significantly higher in AS patients in comparison with healthy controls (17–19).

The unfolded protein response (UPR) is a cellular mechanism implicated in AS pathogenesis (20). Inositol-requiring enzyme 1 (IRE1α), encoded by the AS risk gene ERN1, is an endoplasmic reticulum (ER) stress sensor and UPR initiator (21, 22). IRE1α is activated under ER stress by self-dimerization and phosphorylation, activated IRE1α then cleaves the mRNA of the inactive transcription factor X-box binding protein 1 (XBP1) to generate the mature mRNA that encodes the cleaved XBP-1 (XBP1s) protein, thus upregulating expression of UPR target genes involved in protein folding, processing, and degradation (23, 24). The IRE1α /XBP1 pathway has been shown to be important for the activation of several types of immune cells including macrophages. It has been proposed that the IRE1α /XBP1 pathway is required for toll-like receptor (TLR)-mediated production of pro-inflammatory cytokines IL-6 and TNF-α in macrophages (25).

Why myeloid cells are hyper-responsive to stimulation in AS is unclear. We here provide evidence that a subset of monocytes from AS blood has a strong TI signature and is the main source of pro-inflammatory cytokines in response to either LPS or T-cell stimulation. We show T cell-activated monocytes act back to support T helper (Th17) cell expansion. We identify a population of monocytes in AS joints that closely resembles T cell-activated monocytes, and identify the product of the AS risk gene IRE1α as a (novel) therapeutic target within this pathway.

## RESULTS

### Identification of a subset of monocytes with TI features in blood from AS patients

To identify the molecular and cellular cause of the monocyte hyper-activation in AS, we utilized 10X single-cell RNA sequencing (scRNA-seq) to profile the cellular heterogeneity and transcriptome of peripheral blood mononuclear cells (PBMCs) from 3 ankylosing spondylitis (AS) patients following lipopolysaccharides (LPS) stimulation for 16 hours (Figure S1A). Following the annotation of major immune cell types (Figure S1B and C), 3358 monocytes were acquired and re-clustered (Figure S1D and Figure 1A). In addition to classical and non-classical monocytes, we identified a subset of monocyte expressing IL1B and TNF mRNA in the resting state (Figure 1B) and thought they might have previously experienced microbial stimulation. To test if these cells are “trained” monocytes, we used a set of TI signature genes defined by the transcriptome of trained monocytes induced in vivo through BCG vaccination of healthy donors (2). We found a strong enrichment of TI signature genes in this cluster (Figure 1C-E). Importantly, cells in the trained cluster expressed higher levels of IL1B, TNF and IL6 in response to LPS stimulation (Figure 1F), demonstrating the key functional feature of TI in myeloid cells.

**Figure 1.**
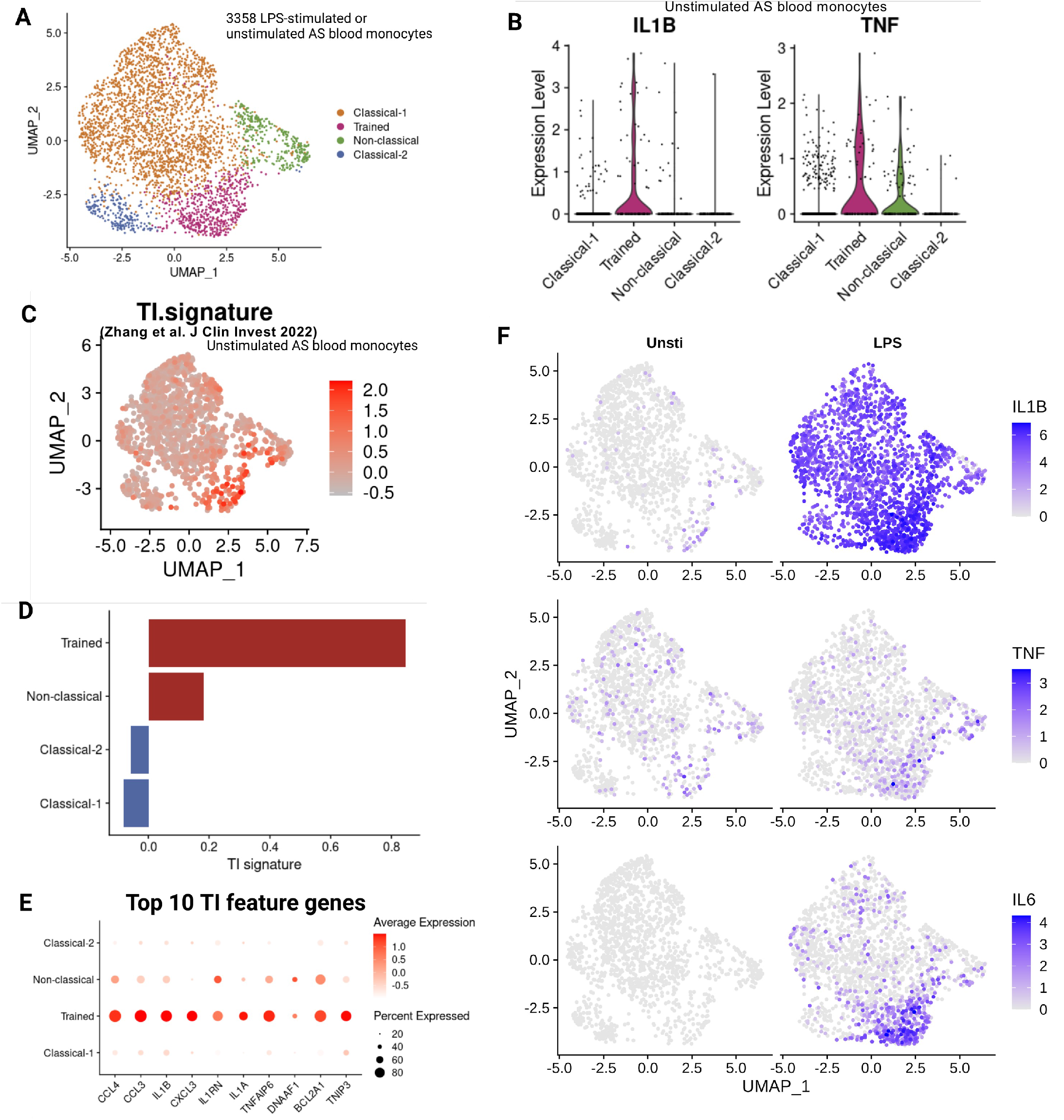
A monocyte subset in AS blood exhibits features of trained immunity. (A) UMAP visualization of transcriptionally distinct populations of monocytes extracted from unstimulated and LPS-stimulated PBMCs. (B) Expression of IL1B and TNF in different monocyte subsets under the unstimulated condition. (C) UMAP visualization showing the trained immunity signature score in unstimulated AS monocytes at the single cell level. (D) Trained immunity signature score of AS unstimulated monocytes at the cluster level. (E) Expression of top 10 trained immunity signature genes by different monocyte subsets (unstimulated). (F) Expression of IL1B, TNF and IL6 projected onto UMAP of monocytes from unstimulated and LPS-stimulated conditions.

### Activated T cells induce cytokine production by blood monocytes from AS patients

Joint inflammation in AS is largely sterile thus the enhanced cytokine production to LPS stimulation in trained monocyte is less likely a major pathological mechanism in AS. Since activated T-cells have been shown to induce IL-1β secretion by myeloid cells in murine models (26), and that T-cell activation is a key feature of AS pathology, we asked if monocytes can be activated by T cells. To this end, we performed intracellular cytokine staining (ICS) for monocytes using PBMCs from 13 AS patients stimulated with LPS or TAB. We gated on monocytes (Figure S2) and observed the induction of IL-1β, TNF-α and IL-6 by both LPS and TAB stimulations (Figure 2A). Of note, TAB induced a level of cytokine production either comparable to (IL-1β), higher (TNF-α) or lower (IL-6) than that of LPS. Since ICS was unable to measure the mature form of IL-1β or quantify IL-23 (very low percentage in ICS), we utilized enzyme-linked immunosorbent assay (ELISA) to confirm the secretion of IL-1β and IL-23 in the culture supernatant following TAB stimulation (Figure 2B). To completely exclude the potential secretion of cytokines from other cells in PBMCs, we carried out a similar experiment using isolated monocytes and CD3+ T cells from the same donors with AS and observed IL-1β, IL-23 and IL-6 production exclusively from T cell/monocyte coculture but not in T-cell alone conditions (Figure 2C). As expected, TNF-α, known to be produced by activated T-cells, was detected in both conditions. Taken together, these data reveal a key mechanism of T cell-instructed monocyte activation leading to proinflammatory cytokine production.

**Figure 2.**
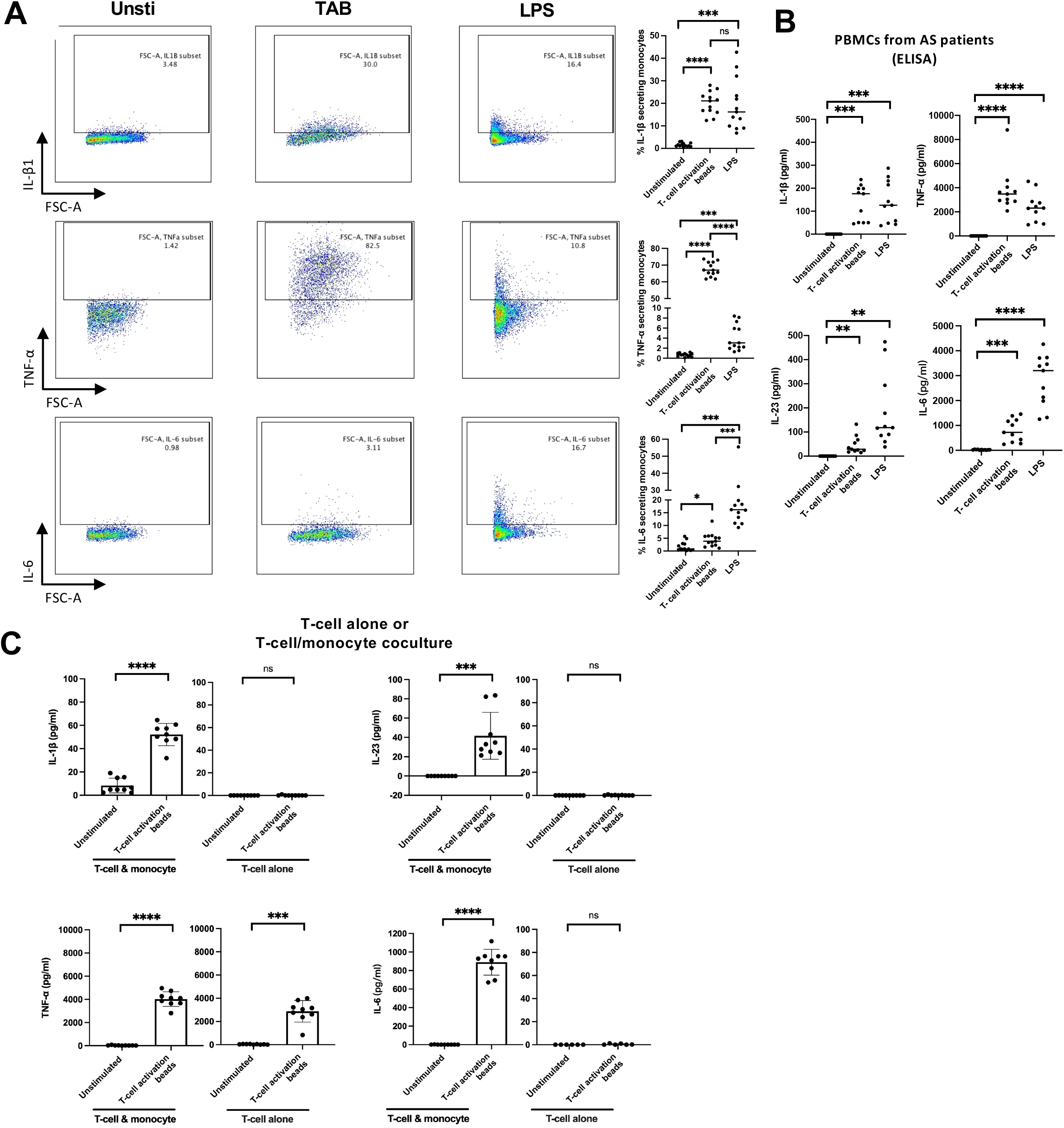
T-cell activation induces cytokine production by Ankylosing Spondylitis monocytes. (A) Representative flow cytometry plots and summary graphs of cytokine-producing CD14+ monocytes from whole PBMCs. (B) Cytokines secreted to supernatant from cultured PBMCs. (C) Isolated CD3+ T-cells from PBMCs from AS patients (n=9) were cultured alone or co-cultured with autologous monocytes in the absence/presence of TAB for 16 hours. The levels of cytokines IL-1β, IL-23 and TNF-α in the culture supernatant were measured using ELISA. The P- value was assessed using the paired two-tailed Student’s t-test (* P≤ 0.05; ** P≤ 0.01; *** P≤ 0.001).

### Trained monocytes are hyper-responsive to T cell-induced activation

We next asked if cells in the trained monocyte cluster (identified in Figure 1) are more responsive to stimulation by T cells. Due to the lack of protein markers for these cells, we used scRNA-seq to profile unstimulated and T cell activated beads (TAB)-stimulated PBMCs using blood cells from 3 AS patients (Figure S3A-C). As expected, genes associated with T cell activation (IFNG and IL2) were induced by TAB in the T cell cluster (Figure S3D). We then extracted monocytes and identified 5 clusters including the subset containing the trained monocytes (Figure 3A). Notably, the higher levels of TNF, IL6 and IL23A were induced by T-cell stimulation in the trained monocyte cluster (Figure 3B).

**Figure 3.**
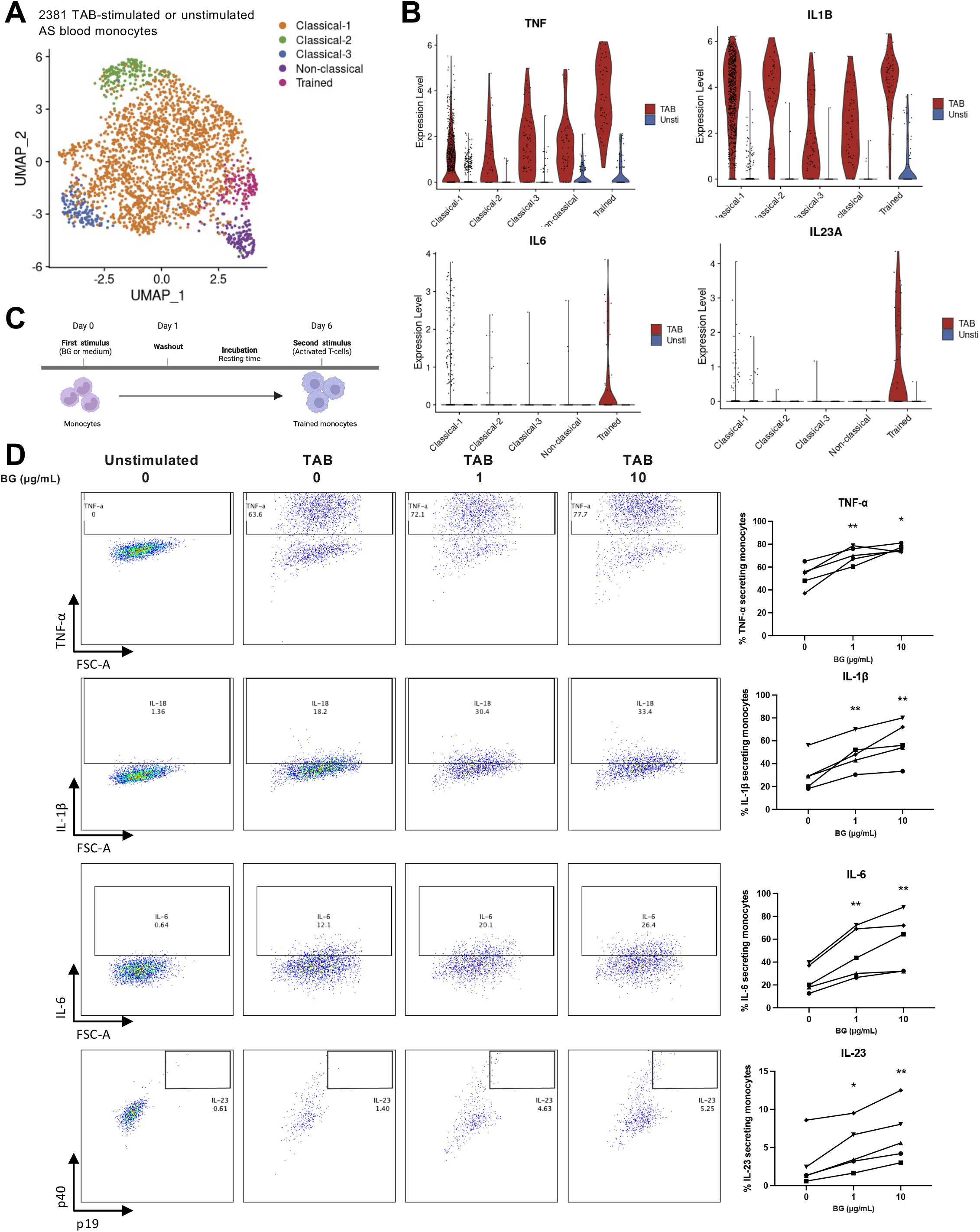
Trained monocytes are hyper-responsive to T-cell-induced activation and act back to T cells to support Th17 response. (A) UMAP visualization of transcriptionally distinct populations of monocytes integrated from unstimulated and TAB-stimulated conditions. (B) Expression of IL1B, TNF, IL6 and IL23A in different monocyte subsets. (C) Experimental schematic of the in vitro β-glucan-induced monocyte training model. (D) Representative flow cytometry plots and summary graphs of cytokine-producing β-glucan-trained monocytes from healthy donors (n=5). The P-value was assessed using the paired two-tailed Student’s t-test (* P≤ 0.05; ** P≤ 0.01; *** P≤ 0.001).

To test if the hyper-response to T cell stimulus is a common feature by TI, we utilized the well-established and widely-used β-glucan-mediated assay to generate trained monocyte in vitro using isolated monocytes from healthy donors (Figure 3C) (3, 5, 7). Following the stimulation using autologous activated T cells, the β-glucan-trained cells produced significantly higher levels of TNF-α, IL-6, IL-1β and IL-23 (Figure 3D). These data show for the first time that, in addition to their enhanced response to microbial stimulus such as LPS, trained monocytes can be hyper-reactive to T cell-induced activation.

### T cell-induced cytokine secretion by monocytes supports Th17 expansion in AS

Th17 cells play a key role in AS pathology and are known to be driven by IL-1β and IL-23, both expressed by monocytes following the stimulation using T cells (Figure 2B and Figure 3B). We hypothesized that T cell-activated monocytes could feed back to T cells to support Th17 expansion via cytokine secretion. To test this hypothesis, we examined the effect of monocyte depletion on the Th17 response in AS PBMCs stimulated using the neutral T cell expansion condition (TAB and IL-2). The percentage of IL-17A-producing CD4+ T cells from patients with AS was significantly downregulated in monocyte-depleted PBMCs compared with whole PBMCs (Figure 4A, gating strategy shown in Figure S4). Strikingly, the Th17-enhancing effect of exogenous IL-1β and IL-23 was only observed in monocyte-depleted PBMCs and not whole PBMCs from AS patients (Figure 4B and C), showing that T cell-induced monocytes cytokine secretion are sufficient to enhance Th17 responses.

**Figure 4.**
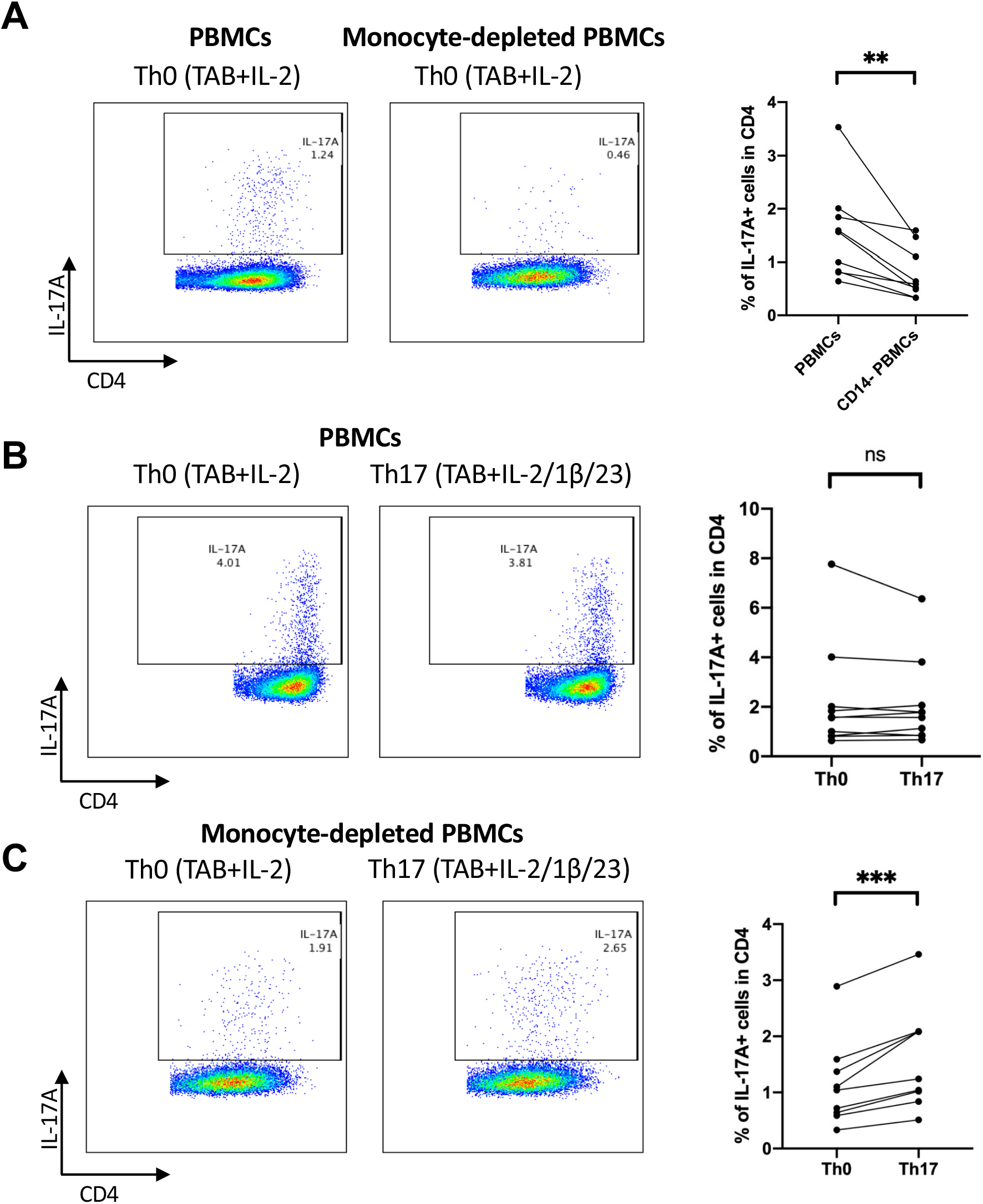
T cell-induced activation of monocytes act back to T cells to support Th17 response in AS. (A) Representative flow cytometry plots showing the percentage of IL-17A- producing CD4+ T cells within the whole PBMCs or monocyte-depleted PBMCs from AS patients (n=9) after 6 days in vitro culture under Th0 condition (left). A summary graph is shown on the right. (B) Representative flow cytometry plots of IL-17A-producing CD4+ T cells from the whole PBMCs from AS patients (n=9) after in vitro culture under Th0 (TAB+IL-2) or Th17 (TAB+IL-2/IL-1β/IL-23) polarizing conditions for 6 days (left). A summary graph is shown on the right. (C) Representative flow cytometry plots of IL-17A-producing CD4+ T cells from the monocyte-depleted PBMCs from AS patients (n=9) after in vitro culture under Th0 or Th17 polarizing conditions for 6 days (left). A summary graph is shown on the right. The P-value was assessed using the paired two-tailed Student’s t-test (* P≤ 0.05; ** P≤ 0.01; *** P≤ 0.001).

### CD14+ myeloid cells in AS synovial fluid are transcriptionally similar to T cell-activated trained monocytes

Next, we asked if trained monocytes that we found in blood is relevant to the pathology of inflammatory synovium in patients with AS. To this end, we examined the expression profile of CD14+ myeloid cells from the synovial fluid (SF) of 8 Ankylosing Spondylitis (AS) patients and their corresponding blood. A total of 16952 CD14+ myeloid cells with high-quality scRNA-seq transcriptomic profiles were obtained (Figures 5A and B). We identified six monocyte clusters but decided to focus on clusters C0-2 as the remaining clusters were scarce in both blood and SF (Figure 5C). CD16+ non-classical monocytes (cluster C0) and Calprotectin+ cluster C1 were more enriched in SF and blood respectively. Cluster C2, dominated by SF-sourced monocytes, was the main source of IL1B and TNF (Figure 5D-E), key pro-inflammatory cytokines in AS pathology. Multiple additional pro-inflammatory cytokines and chemokines were highly enriched in cluster C2 (Figure 5F). The upregulation of IL1B and IL23A, as well as CCL20 (a known Th17 recruiter through CCR6), suggests that cluster C2 could enhance both enrichment and expansion of Th17 cells.

**Figure 5.**
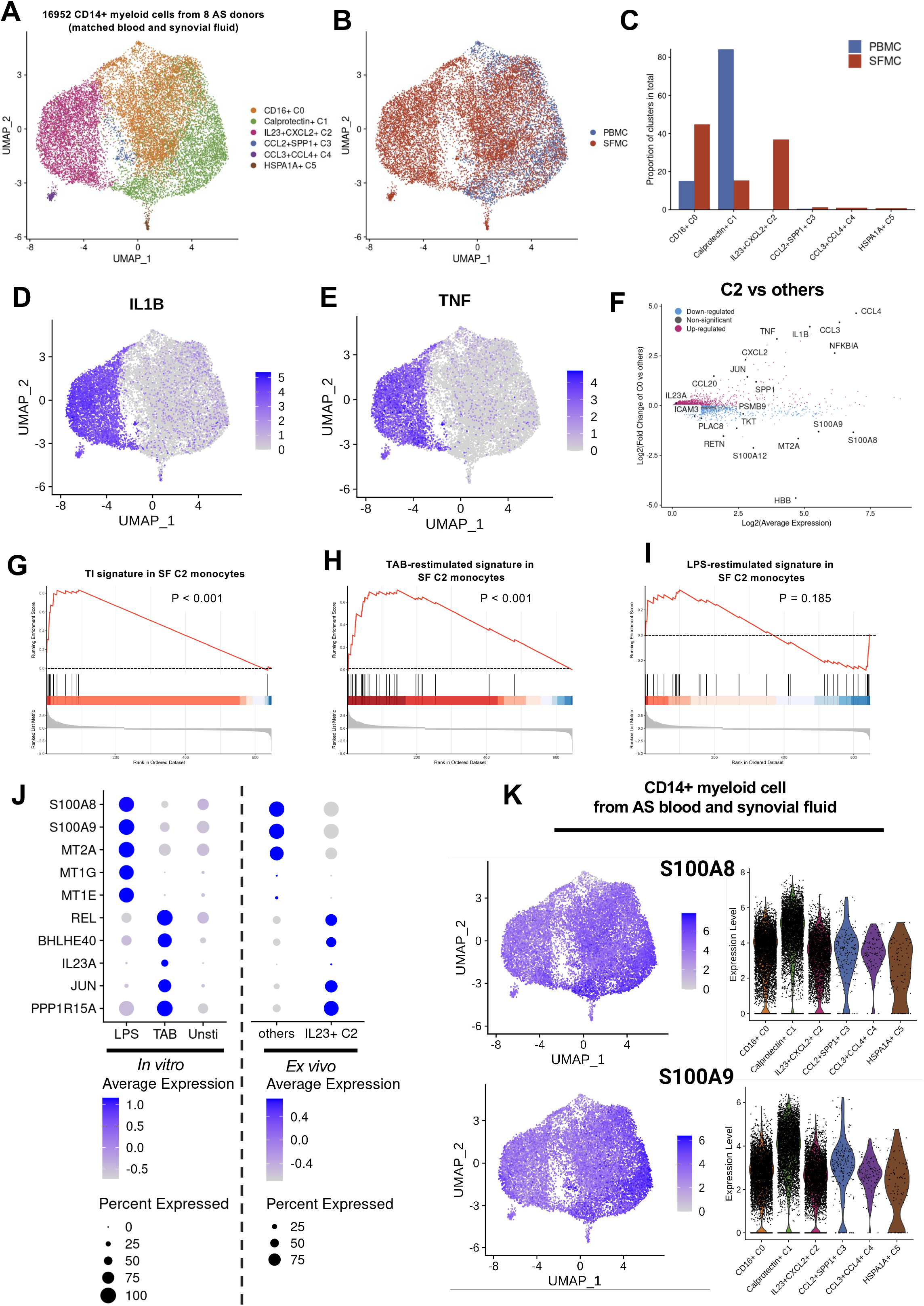
CD14+ myeloid cells in AS synovial fluid are transcriptionally similar to T cell- stimulated trained monocytes. (A) UMAP visualization of transcriptionally distinct populations of CD14+ myeloid cells from 8 paired AS blood and synovial fluid samples. (B) UMAP visualization of CD14+ myeloid cell from blood and synovial fluids, colored by source. (C) Frequency of cells from various clusters in PBMCs and SFMCs. (D) UMAP visualization of IL1B expression by CD14+ myeloid cells in AS synovial fluid. (E) UMAP visualization of TNF expression by CD14+ myeloid cells in AS synovial fluid. (F) MA plot of differentially expressed genes in IL23+CXCL2+ C2 cells versus the rest (FDR <0.05). The average expression levels are shown for C2 when the fold change is greater than 0, and for the others when the fold change is less than 0. GSEA analysis was carried out for SF C2 cluster to test of signature genes in trained monocyte (G), T cell-activated trained monocytes from Figure 3A and B (H) or LPS-activated trained monocytes from Figure 1A and B (I). Further details are described in the method section. (J) Expression of a set of representative genes by monocytes from in vitro and ex vivo experiments. (K) Expression of S100A8 and S100A9 by distinct monocyte clusters from AS blood and joints.

We then checked if TI is relevant to cluster C2 using the same approach used for Figure 1C-E and observed a significant enrichment of TI signature genes (Figure 5G). Notably, Cluster C2 was only enriched for the T cell-activated trained monocyte gene signature not LPS-activated (Figure 5 H and I). Figure 5J shows that genes specifically induced by LPS (S100A8, S100A9, MT2A, MT1G and MT1E) in trained monocytes were expressed at lower levels in the IL23+ cluster C2 largely formed by cells from synovium fluid (Figure 5C). By contrast, many genes such as REL, BHLHE40, IL23A, JUN and PPP1R15A were enriched in both T cell-activated trained monocytes and in the IL23+ cluster C2. Notably, S100A8 and S100A9 were more abundant in cluster C1 (Figure 5K) which is mostly composed of cells from blood (Figure 5C), further supporting that T cells activated trained monocytes in inflamed synovium.

### Activated T-cells are present in AS synovial fluid and are composed of both CXCR6+ and CXCR6-T-cells

We next looked for evidence of T-cell activation in SF using T cells from the matched AS synovial and blood datasets. A total of 12629 T-cells were acquired revealing 8 major populations (Figure 6A and B, Figure S5). Clear enrichment of PDCD1+IFNG+ activated T-cells was observed in memory CD4+ and CD8+ T-cells from SFMCs compared to those from PBMCs (Figure 6C). Notably, these activated T-cells were composed of both CXCR6+ tissue resident and CXCR6-circulating T-cells (Figure 6D), suggesting that these effector cells could have been either locally generated in the joints or migrated from other organs (such as the gut). Overall, these findings suggest that activated T-cells were present in AS SF that could provide stimulation to CD14+ myeloid cells in SF.

**Figure 6.**
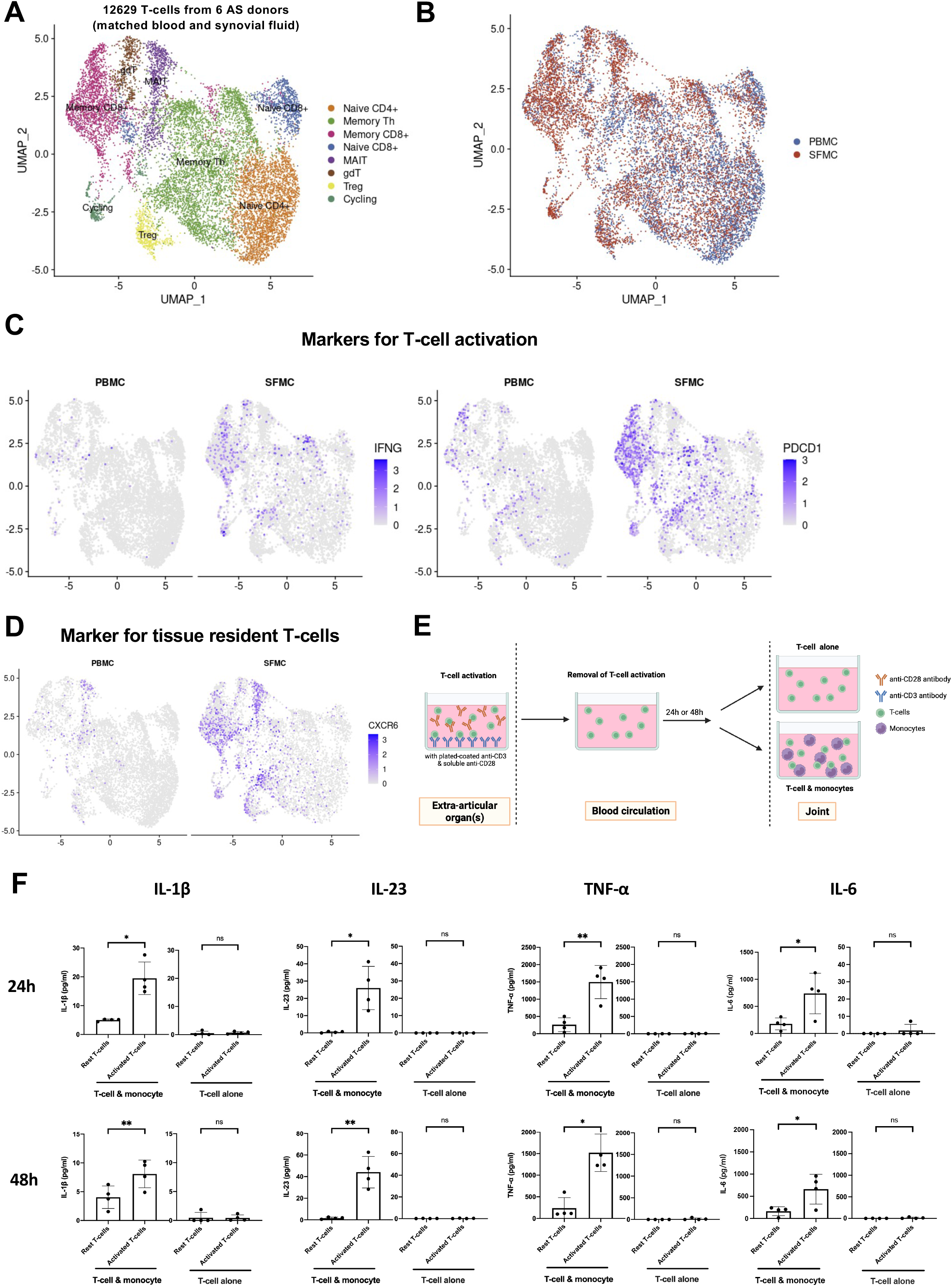
Activated T-cells are present in synovial fluid from patients with AS. (A) UMAP visualization of T cells subsets from 6 pairs of matched blood and synovial fluid samples from AS patients. (B) UMAP visualization of T cells from blood and synovial fluid, colored by the source of cells. (C) Expression of T cell activation marker genes (IFNG and PDCD1) projected onto a UMAP of T cells from AS blood and synovial fluid. (D) Expression of CXCR6 projected onto a UMAP of T cells from AS blood and synovial fluid. (E) Experimental schematic of the in vitro assay modelling the activation of monocytes in joints by T-cells activated in extra-articular organs. CD3+ T-cells (green) were isolated from PBMCs of AS patients and cultured overnight in the presence or absence of anti-CD3 (blue, plate-coated) and anti-CD28 (yellow, soluble) antibodies. Unstimulated and stimulated CD3+ T-cells were then cultured in the fresh culture medium for 24 or 48 hours before being co-cultured with the monocytes (purple) from the same donors for 16 hours. (F) The levels of cytokines IL-1β, IL-23, TNF-α and IL-6 in the culture supernatant were then measured by ELISA (n=4).

Multiple pieces of evidence have supported the enrichment of gut-derived T-cells in inflamed AS joints (27, 28). Thus, the CXCR6-PDCD1+IFNG+ activated T-cells that we observed in AS SF were likely generated in the gut. To test whether T-cells activated in extra-articular organ(s) could maintain their ability to activate monocytes following migration to the joints, we designed an in vitro assay to model this biological process (Figure 6E). We found that T cells pre-activated either 24 or 48 hours before co-culture preserved their ability to induce monocytes to produce pro-inflammatory cytokines IL-1β, IL-23, IL6 and TNF-α (Figure 6F). This finding, together with data shown in Figure 2, suggests that T-cells either locally activated in synovium or pre-activated e.g. in the gut are capable of inducing monocyte activation in joints.

### T cell-activated trained monocytes express high levels of AS-associated risk genes and are regulated by the AS risk gene ERN1

In order to identify key regulators determining the function of trained monocyte in AS, we next looked for the expression of AS risk genes identified in Genome-wide association studies (GWASs). We found 8 AS risk genes whose expression was enriched in trained monocytes activated by T cells (Figure 7A). The ERN1 gene was particularly interesting as its protein product IRE1α is a key sensor for endoplasmic reticulum (ER) stress, a key hypothesized cause of AS and myeloid cell activation (25, 29). In addition, using a publicly available dataset (ImmuNexUT), we found that the AS risk allele (G) is associated with an increased ERN1 expression level in human primary monocytes (Figure 7B). These pieces of evidence encouraged us to hypothesize that ERN1 might contribute to T cell-instructed activation of trained monocytes. We used siRNA to suppress IRE1α expression in isolated primary monocytes from patients with AS and observed significant reduction in the production of IL-1β and IL-23 (Figure 7C and D). In line with this, we found that 4μ8C, a widely used inhibitor specific for IRE1α, significantly reduced the production of IL-1β and IL-23 using the same assay (Figure 7E). Taken together, these results reveal a key role of IRE1α in the production of proinflammatory cytokines in T cell-stimulated trained monocytes.

**Figure 7.**
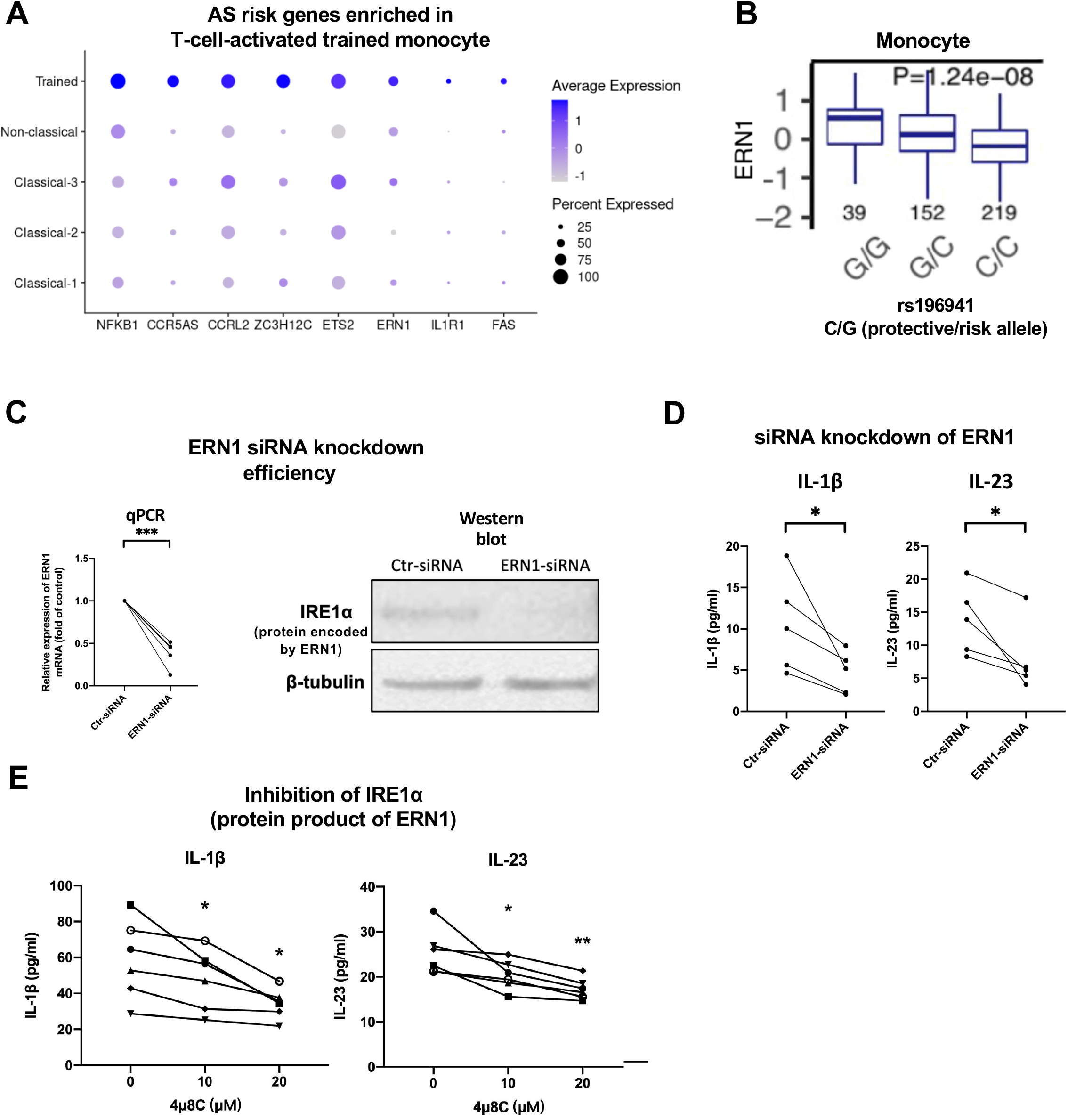
T cell-activated trained monocytes express high levels of AS-associated risk genes and are regulated by the AS risk gene ERN1. (A) Expression of AS risk genes enriched in T cell-activated trained monocytes. (B) A higher expression level of ERN1 in monocytes is associated with AS risk allele G at SNP rs196941 (using publicly available data from ImmuNexUT). (C) Efficient knockdown of ERN1 at mRNA and protein (IRE1α) levels. (D) Isolated monocytes from AS patient blood (n=5) were transfected with ERN1 siRNA or non-targeting control siRNA before the co-culture with autologous CD3+ T cells and T cell activation beads for 16 hours. The levels of IL-1β and IL-23 in the culture supernatant were measured by ELISA. (E) Isolated monocytes from AS patient blood (n=6) were treated with an IRE1α inhibitor (4u8C) for 17 h. The 4u8C was then washed away before 4 h co-culture with autologous CD14-depleted PBMCs and T cell activation beads. P-values assessed using paired two-tailed hStudent’s t-test (* P≤ 0.05; ** P≤ 0.01; *** P≤ 0.001).

## DISCUSSION

In this study, we show that trained immunity (TI) is present in a subset of peripheral monocytes in patients with Ankylosing Spondylitis (AS). These “trained” monocytes are enriched for TI signatures and are hyper-responsive to LPS stimulation, a key functional feature of TI. Interestingly, we found the “trained” monocytes to exhibit enhanced cytokine response to stimulation induced by activated T cells and confirm this finding using the classic β-glucan monocyte TI training assay. To our knowledge, this feature of TI monocytes has not been reported before. This may not only contribute to AS pathogenesis, but could provide important insights into other scenarios related to both TI and T cell responses, such as vaccination and Alzheimer’s disease.

It would be interesting to check whether the subset of trained monocytes that we identified in AS is present in other forms of inflammatory arthritis. Monocyte hyper- activation has been reported in other forms of inflammatory arthritis, such as Rheumatoid Arthritis, Psoriatic Arthritis and Juvenile Idiopathic Arthritis (JIA) (30–32). Investigation of TI in these fields could be of value in advancing our knowledge in pathological mechanisms.

We also show that cytokines produced by T cell-stimulated monocytes act back on T cells to enhance Th17 responses, a major component of AS pathology. The expansion of Th17 cells in PBMCs from AS patients was reduced by monocyte depletion and rescued by the addition of exogenous Th17 skewing cytokines (IL-1β and IL-23). This suggests that monocytes in AS have the capacity to provide sufficient Th17-supporting cytokines in the absence of pathogen-associated molecular patterns (PAMPs) such as LPS. This mechanism of monocyte activation is likely relevant to the sterile joint inflammation seen in AS and related inflammatory arthritides.

Notably, the proinflammatory CD14+ myeloid cells identified here in the joints of patients with AS are enriched for the gene signature of T cell-stimulated rather than LPS-stimulated trained monocytes. In line with this, and in contrast to the lack of evidence supporting an abundance of PAMPs in AS joints, we observed enrichment of PD1+IFN-γ+ T-cells in AS synovial fluid. Our data thus suggest that T-cell-induced monocyte activation is actively occurring in inflamed AS joints and likely contributes to pathology. We also show that T cells activated 24 to 48 hours previously are capable of activating monocytes - they would thus have time to migrate to the joints following activation at other sites (eg gut). This data is consistent with models of either joint or distant T cell activation with subsequent effector function in the joint. Indeed, we here provide evidence that monocyte activation may indeed be a key part of this effector function.

Interestingly, around half of the activated T-cells we observed in AS SF were CD8+, supporting previous studies proposing that antigen-experienced CD8+ T cells travel from gut to joint and then undergo local expansion (27, 28). In addition, a recent study reports the presence of HLA-B*27-restricted peptides that activate T cell receptors present in AS joint CD8+ T cells (33). Our data along with these published findings imply a role of (potentially gut-derived) HLA-B*27-restricted CD8+ T-cells in the activation of CD14+ myeloid cells in joints, which leads to the upregulation of proinflammatory cytokines such as IL-1β and IL-23. These Th17-supporting cytokines could be upstream of IL-17 production in the synovium. Future work needs to be carried out to test this hypothesis.

Drug targets with genetic linkage are more likely to reach late-stage clinical trials (34, 35). We found that ERN1, an AS risk gene, was enriched in T cell-activated trained monocytes. Silencing of ERN1 or inhibiting the function of its gene product (IRE1α protein) both reduced the production of IL-1β and IL-23 by T cell-activated monocytes from patients with AS. Thus, the inhibition of IRE1α or its related pathway is an attractive approach for AS treatment.

The enhancement of unfolded protein response (UPR) is a key hypothesized cause of AS, largely driven the study of HLA-B*27 function in AS. Considering the key role of IRE1α in UPR, it is plausible to predict that HLA-B*27 would be involved in the function of trained monocytes in AS. Future work needs to be carried out to test this hypothesis.

Gut inflammation and leakage have been found in a proportion of patients with AS, accompanied by the upregulation of IL-1β and IL-23 in inflamed gut tissue (36–38). Accordingly, a model in which monocytes are activated by bacteria in the gut or by bacterial derivatives in the bloodstream has been considered (36). We propose that the trained monocytes that we identified in AS blood were likely “educated” in the gut and subsequently re-activated by T cells in joints.

In summary, we here identify the presence of trained immunity in monocytes from patients with AS and show for the first time that trained monocytes are hyper-responsive to T cell stimulation. Additionally, our data reveal the presence of a pro-inflammatory circuit initiated by T cell-induced activation of trained monocytes which then can act back to enhance Th17 responses. These findings thus advance our knowledge of trained immunity and AS pathology and offer therapeutic potential.

## METHODS

### Patient and Control Recruitment

Peripheral blood was collected from patients attending the Oxford University Hospitals National Health Service Foundation Trust (OUH), with appropriate consent and ethical approval (Ethics reference number 06/Q1606/139, National Health Service, Health Research Authority, South Central - Oxford C Research Ethics Committee). All patients included in this study met the criteria of the Assessment of Spondyloarthritis International Society (ASAS) for AxSpA (which we henceforth term AS). Fifteen of the patients underwent biological therapy (adalimumab, etanercept, secukinumab or Golimumab), and twenty-eight patients were treated with non-steroidal anti-inflammatory drugs (Ibuprofen, etoricoxib, naproxen, Meloxicam). The demographics of AS patients recruited for this study are shown in **Table S1**.

### Cell isolation and culture

Human PBMCs were obtained from the whole blood of AS patients by density gradient centrifugation using Histopaque-1077 (Sigma-Aldrich). Primary human monocytes and CD3+ T cells were isolated from PBMCs using CD14 MicroBeads™ and pan T cell isolation kit (Miltenyi Biotec), respectively. Human PBMCs, CD14+ monocytes or CD14-depleted PBMCs were cultured in RPMI 1640 medium (Sigma) supplemented with 10% fetal bovine serum (FBS) and 1% GlutaMAX (Gibco) at 37°C in a humidified atmosphere containing 5% carbon dioxide.

### Monocyte activation

For monocyte stimulation assays, isolated 1×10^6^ PBMC were cultured in 200 μl medium in non-tissue culture-treated 96-well round bottom plates (Corning) and stimulated with 200 ng/ml LPS (Enzo Bioscience) or anti-CD2/CD3/CD28-coated T-cell activation/expansion beads (Miltenyi Biotec) for 16h. To exclude the involvement of non-monocytes in PBMCs, monocytes and T-cells were isolated from PBMCs (90- 95% purity) and co-cultured in the presence of TAB for 16 hours. For inhibition assays, 0.2 x 10^6^ CD14+ monocytes were pre-treated with the IRE1α inhibitor 4μ8C (0, 10 and 20uM) for 17 h. The 4μ8C was then washed away before the 4 h co-culture with autologous CD14-depleted PBMCs and anti-CD2/CD3/CD28-coated beads. For siRNA experiment, siRNA-transfected CD14+ monocytes were stimulated with isolated autologous CD3+ T-cell and anti-CD2/CD3/CD28-coated beads.

### Monocyte β-glucan training assay

0.2 x 10^6^ isolated CD14+ monocytes were incubated with 0, 1 or 10 μg/mL of β-glucan (InvivoGen) for 24 hours in 24 well plates. 24 hours later, β-glucan was removed through washing and the use of cell strainer (Corning, 352235). Monocytes were then cultured in RPMI supplemented with 10% pooled human serum, 2mM Glutamax (GIBCO) and 1 mM pyruvate (GIBCO) for additional 5 days before the stimulation by activated T cells. For each well, 0.4 x 10^6^ isolated T cells from the same donor and 0.4 x 10^6^ T cell activation beads (Miltenyi Biotec, anti-CD2/3/28) were added. 4 hours post T cell activation, BFA was added. Cells were harvested 16 hours after T cell stimulation for cytokine staining using ICS.

### Preparation of samples and libraries for scRNA-seq

Following stimulation, PBMCs were then washed in PBS with 0.1% BSA. Blood samples from donors AS1802, AS1830, AS2311 were used in this experiment. For AS1802, we loaded 20,000 cells from each condition into a channel of Chromium Next GEM Chip K of the 10x Genomics platform. For AS1830 and AS2311, we multiplexed cells from the same conditions at a 1:1 ratio using Total-seq-C hashtag antibodies (C0251 and C0252), and loaded 20,000 multiplexed samples from each reaction onto the chip separately. Single-cell RNA-sequencing libraries were generated using the Chromium Next GEM Single Cell 5’ v2 kit according to the manufacturer’s instructions. The resulting DNA libraries were sequenced at >50,000 reads/cell on a Novoseq 6000 using PE150 mode. For two pairs of AS blood and synovial fluid samples, we first isolated PBMCs and SFMCs and then performed fluorescence-activated cell sorting to enrich for CD3- live cells. We used the same library construction and sequencing strategies as in the in vitro stimulation experiment, except we used the Chromium Next GEM Single Cell 5’ v1 kit instead of v2. For the additional six donors, we profiled whole PBMCs and SFMCs using methods that have been described previously (39).

### Computational analysis of scRNA-seq data

The paired reads obtained were mapped to the hg38 reference genome to generate gene expression matrices using CellRanger v7.0.0. The raw matrices were then analyzed using the Seurat R package (v4.0.5). Cells with low-quality profiles were excluded based on the number of detected genes, the percentage of mitochondrial RNA among total UMIs, and the total number of UMIs. The raw read counts were normalized using the NormalizeData function, and variable genes were identified for each sample. Data across different conditions were then integrated using the SCTtransform function. To select candidate anchor genes, genes were first selected using the SelectIntegrationFeatures function, and TCR genes were then further excluded.

To identify cell types, we performed principal component analysis (PCA) on integrated data with the RunPCA function to generate principal components (PCs) used for dimensionality reduction. PCs representing >90% variance were used for Uniform Manifold Approximation and Projection (UMAP) dimensionality reduction. The FindNeighbors and FindCluster functions were used to cluster cells based on global transcriptional profile. Clusters in the 2-dimensional UMAP were used to identify cell types based on marker genes. When re-clustering CD14+ myeloids, CD14+ populations were extracted, and dendritic cells were further excluded based on the expression of FLT3, CLEC9A, and CLEC10A. Pure CD14+ myeloids were then split by donors and re-integrated to capture the variance within this population. For differential expression analysis, the FindMarkers function was used to test the normalized data with the default Wald test method. Unless specified, default parameters were used for each function.

To calculate the TI (trained immunity) signature scores for profiled cells, TI signature genes were defined as genes up-regulated (adjusted p-value < 0.05 and log2 fold change > 0.3) in blood monocytes 3 months post BCG vaccination compared to pre-vaccination (2). The normalized and scaled counts of TI signature genes of cells were summed and divided by the number of genes used. The average TI signature score of cells within a certain cluster was used for the cluster-level scoring.

### T cell expansion assays

PBMCs and CD14-depleted PBMCs were cultured in 96-well u-bottom plates at a density of 5 × 10^5^ per well and stimulated with anti-CD2/CD3/CD28-coated T-cell activation/expansion beads (Miltenyi Biotec) in the presence of IL-2 (20 ng/ml, PeproTech) for 6 days. For Th17 cell cultures, recombinant human IL-23 (20 ng/mL, PeproTech) and IL-1β (20 ng/mL, PeproTech) were added at the initiation of culture. For Th0 generation, cells were stimulated as above in absence of IL-1β and IL-23. On day 3, half of the culture medium was replaced with the fresh medium containing the same concentration of IL-2, IL-23 and IL-1β. On day 6, cells were harvested for flow cytometry staining.

### siRNA silencing of ERN1 gene in primary monocytes

Primary monocytes were cultured in 24-well plates and then transfected with control non-targeting siRNA (Dharmacon) or siRNA targeting ERN1 (Dharmacon) using HiPerFect reagent (QIAGEN) following the instructions provided by the manufacturer. After 3 days, the cells were collected for subsequent experiments. The knockdown efficiency was determined by quantitative PCR (qPCR) and western blotting. For qPCR, primary monocytes were pelleted and resuspended in TRIzol. Total RNA was extracted using the Direct-zol RNA MiniPrep Kit according to the manufacturer’s instructions (Zymo Research, R2052). The RNA was then reverse transcribed into cDNA using the High-Capacity cDNA Reverse Transcription Kit (Applied Biosystems, 4368813). Using Taqman Gene Expression Assay probe Hs00980095_m1 for ERN1 and Hs1060665_g1 for ACTB (Applied Biosystems), quantitative real-time PCR was performed with an Applied Biosystems Prism 7500 Fast Sequence Detection System using the TaqMan fast universal PCR master mix reagents (Applied Biosystems, 4352042). The mRNA expression levels were analysed using the 2–ΔΔCt method.

For western blotting, primary monocytes were lysed using RIPA buffer supplemented with protease inhibitors (Sigma). Protein concentrations were quantified by bicinchoninic acid (BCA) assay. Total proteins (18 μg) were electrophoresed through a 12% sodium dodecyl sulfate (SDS)–polyacrylamide gels, transferred to the polyvinylidene difluoride membranes and analyzed by immunoblotting. The primary antibodies used were rabbit monoclonal anti-IRE1α (1:1000 dilution; Cell Signalling; 14C10) and mouse anti-α-Tubulin (1:1000 dilution; Cell Signalling; DM1A). The secondary antibodies used were goat anti-rabbit IgG (1:35000, Bethyl Laboratories, A120-201P) or goat anti-mouse IgG (1:25000, Bethyl Laboratories, A90-516P).

### Intracellular cytokine staining (ICS)

T cells were stimulated with 100ng/mL PMA (Sigma-Aldrich) and 1ug/mL Ionomycin (Sigma-Aldrich) for 2.5 h followed by 2.5 h incubation with 5 μg/ml Brefeldin-A (Sigma-Aldrich) before intracellular cytokine staining. The following FACS anti-human antibodies were used: CD3 (clone OKT3; Biolegend); CD4 (clone RPA-T4; BioLegend) and IL-17A (clone eBio64DEC17; eBioscience).

For monocyte experiments, PBMCs were stimulated with either LPS or T cell activation beads for 4 h followed by 12 h incubation with Brefeldin-A (Sigma-Aldrich). Following FACS anti-human antibodies were used for analysis: CD3 (clone OKT3; Biolegend); CD14 (clone M5E2; Biolegend); IL-1 beta (clone CRM56 eBioscience); IL-6 (MQ2-13A5; eBioscience); XBP1s (Q3-695; BD Biosciences) and TNF-a (clone Mab11; BioLegend). To identify living cells and exclude dead cells, samples were stained with LIVE/DEAD™ Fixable Violet Dead Cell Stain Kit (L34955, Invitrogen). For intracellular cytokines and transcription factor staining, cells were fixed and permeabilized in the Transcription Factor Buffer Set (562574, BD Biosciences). Data analysis was performed using FlowJo software.

### Enzyme-linked immunosorbent assays (ELISA)

The concentrations of IL-23, IL-1β, TNF-α and IL-6 in cell culture supernatants were assayed using standard ELISA kits (Invitrogen or R&D) in accordance with the protocol set out by the manufacturer. The Multiskan Ascent ELISA reader was employed to read the ELISA plates at a test wavelength of 450 nm and a background wavelength of 570 nm. Then, the standard curve of ELISA was plotted using Microsoft Excel software, while the corresponding concentrations of samples were calculated using the absorbance value and standard curve.

### Statistics

The level of statistical significance was assessed by paired Student’s t-test. The differences were considered statistically significant at P<0.05*, P<0.01**, P<0.001***. All statical analyses and summarized graphs were performed using GraphPad Prism 8.4.0.

### Study approval

Venous blood and synovial fluid were obtained under protocols approved by the Oxford Research Ethics committee (Ethics reference number 06/Q1606/139).

## Data availability

Raw and processed data of the in vitro stimulation experiment are available under the GEO accession number GSE232131. Data of paired blood and synovial fluids is publicly available through Zenodo (https://doi.org/10.5281/zenodo.7730757).

## Author contributions

JZ, FL, PB and LC designing research studies, JZ, FL, CW, DS, FP, HS, JL, SL, HF, JSL, TJK, LD conducting experiments, JZ, FL, CW, DS, FP, HS, JL, SL, HF, JSL, TJK, LD acquiring data, JZ, FL, CW, DS, FP, HS, JL, SL, HF, JSL, TJK, LD analyzing data, JZ, FL, PB and LC writing the manuscript. JZ and FL are co–first authors. JZ and FL have led wet-lab and dry-lab work for this study respectively and given the co-first authorship as the experimental and bioinformatic findings are equally important to support the main findings in this study. JZ was placed at the first place as she has spent more time on this study than FL.

## Acknowledgements

This work was funded by a Versus Arthritis career development award to LC 22053. P.B. is funded by the National Institute for Health Research (NIHR) Oxford Biomedical Research Centre (BRC). The views expressed are those of the author(s) and not necessarily those of the NHS, the NIHR or the Department of Health. PB FP and DS are funded by a Versus Arthritis award grant number 22252.

## REFERENCES

1. Netea MG, Quintin J, van der Meer JW. Trained immunity: a memory for innate host defense. Cell Host Microbe. 2011;9(5):355–61.

2. Zhang B, Moorlag SJ, Dominguez-Andres J, Bulut Ö, Kilic G, Liu Z, et al. Single-cell RNA sequencing reveals induction of distinct trained-immunity programs in human monocytes. J Clin Invest. 2022;132(7).

3. Quintin J, Saeed S, Martens JHA, Giamarellos-Bourboulis EJ, Ifrim DC, Logie C, et al. Candida albicans infection affords protection against reinfection via functional reprogramming of monocytes. Cell Host Microbe. 2012;12(2):223–32.

4. Joosten SA, van Meijgaarden KE, Arend SM, Prins C, Oftung F, Korsvold GE, et al. Mycobacterial growth inhibition is associated with trained innate immunity. J Clin Invest. 2018;128(5):1837–51.

5. Saeed S, Quintin J, Kerstens HH, Rao NA, Aghajanirefah A, Matarese F, et al. Epigenetic programming of monocyte-to-macrophage differentiation and trained innate immunity. Science. 2014;345(6204):1251086.

6. Hata M, Andriessen E, Hata M, Diaz-Marin R, Fournier F, Crespo-Garcia S, et al. Past history of obesity triggers persistent epigenetic changes in innate immunity and exacerbates neuroinflammation. Science. 2023;379(6627):45–62.

7. Cheng SC, Quintin J, Cramer RA, Shepardson KM, Saeed S, Kumar V, et al. mTOR- and HIF-1α-mediated aerobic glycolysis as metabolic basis for trained immunity. Science. 2014;345(6204):1250684.

8. Christ A, Günther P, Lauterbach MAR, Duewell P, Biswas D, Pelka K, et al. Western Diet Triggers NLRP3-Dependent Innate Immune Reprogramming. Cell. 2018;172(1-2):162–75.e14.

9. Brown MA, Kenna T, Wordsworth BP. Genetics of ankylosing spondylitis--insights into pathogenesis. Nat Rev Rheumatol. 2016;12(2):81–91.

10. Jandus C, Bioley G, Rivals JP, Dudler J, Speiser D, Romero P. Increased numbers of circulating polyfunctional Th17 memory cells in patients with seronegative spondylarthritides. Arthritis Rheum. 2008;58(8):2307–17.

11. Shen H, Goodall JC, Hill Gaston JS. Frequency and phenotype of peripheral blood Th17 cells in ankylosing spondylitis and rheumatoid arthritis. Arthritis Rheum. 2009;60(6):1647–56.

12. Baeten D, Sieper J, Braun J, Baraliakos X, Dougados M, Emery P, et al. Secukinumab, an Interleukin-17A Inhibitor, in Ankylosing Spondylitis. N Engl J Med. 2015;373(26):2534–48.

13. Chung Y, Chang SH, Martinez GJ, Yang XO, Nurieva R, Kang HS, et al. Critical regulation of early Th17 cell differentiation by interleukin-1 signaling. Immunity. 2009;30(4):576–87.

14. McGeachy MJ, Chen Y, Tato CM, Laurence A, Joyce-Shaikh B, Blumenschein WM, et al. The interleukin 23 receptor is essential for the terminal differentiation of interleukin 17-producing effector T helper cells in vivo. Nat Immunol. 2009;10(3):314–24.

15. Wilson NJ, Boniface K, Chan JR, McKenzie BS, Blumenschein WM, Mattson JD, et al. Development, cytokine profile and function of human interleukin 17–producing helper T cells. Nature Immunology. 2007;8(9):950–7.

16. Conrad K, Wu P, Sieper J, Syrbe U. In vivo pre-activation of monocytes in patients with axial spondyloarthritis. Arthritis Res Ther. 2015;17(1):179.

17. Shi H, Chen L, Ridley A, Zaarour N, Brough I, Caucci C, et al. GM-CSF Primes Proinflammatory Monocyte Responses in Ankylosing Spondylitis. Frontiers in Immunology. 2020;11.

18. Zeng L, Lindstrom MJ, Smith JA. Ankylosing spondylitis macrophage production of higher levels of interleukin-23 in response to lipopolysaccharide without induction of a significant unfolded protein response. Arthritis Rheum. 2011;63(12):3807–17.

19. Mei Y, Pan F, Gao J, Ge R, Duan Z, Zeng Z, et al. Increased serum IL-17 and IL-23 in the patient with ankylosing spondylitis. Clin Rheumatol. 2011;30(2):269–73.

20. Navid F, Colbert RA. Causes and consequences of endoplasmic reticulum stress in rheumatic disease. Nat Rev Rheumatol. 2017;13(1):25–40.

21. Schröder M, Kaufman RJ. The mammalian unfolded protein response. Annu Rev Biochem. 2005;74:739–89.

22. Ron D, Walter P. Signal integration in the endoplasmic reticulum unfolded protein response. Nat Rev Mol Cell Biol. 2007;8(7):519–29.

23. Janssens S, Pulendran B, Lambrecht BN. Emerging functions of the unfolded protein response in immunity. Nat Immunol. 2014;15(10):910–9.

24. Tavernier SJ, Osorio F, Vandersarren L, Vetters J, Vanlangenakker N, Van Isterdael G, et al. Regulated IRE1-dependent mRNA decay sets the threshold for dendritic cell survival. Nat Cell Biol. 2017;19(6):698–710.

25. Martinon F, Chen X, Lee AH, Glimcher LH. TLR activation of the transcription factor XBP1 regulates innate immune responses in macrophages. Nat Immunol. 2010;11(5):411–8.

26. Jain A, Irizarry-Caro RA, McDaniel MM, Chawla AS, Carroll KR, Overcast GR, et al. T cells instruct myeloid cells to produce inflammasome-independent IL-1β and cause autoimmunity. Nat Immunol. 2020;21(1):65–74.

27. Qaiyum Z, Gracey E, Yao Y, Inman RD. Integrin and transcriptomic profiles identify a distinctive synovial CD8+ T cell subpopulation in spondyloarthritis. Ann Rheum Dis. 2019;78(11):1566–75.

28. Guggino G, Rizzo A, Mauro D, Macaluso F, Ciccia F. Gut-derived CD8(+) tissue-resident memory T cells are expanded in the peripheral blood and synovia of SpA patients. Ann Rheum Dis. 2021;80(11):e174.

29. Bowness P. HLA-B27. Annu Rev Immunol. 2015;33:29–48.

30. Zhang F, Wei K, Slowikowski K, Fonseka CY, Rao DA, Kelly S, et al. Defining inflammatory cell states in rheumatoid arthritis joint synovial tissues by integrating single-cell transcriptomics and mass cytometry. Nat Immunol. 2019;20(7):928–42.

31. Yager N, Cole S, Lledo Lara A, Maroof A, Penkava F, Knight JC, et al. Ex vivo mass cytometry analysis reveals a profound myeloid proinflammatory signature in psoriatic arthritis synovial fluid. Ann Rheum Dis. 2021;80(12):1559–67.

32. Cepika AM, Banchereau R, Segura E, Ohouo M, Cantarel B, Goller K, et al. A multidimensional blood stimulation assay reveals immune alterations underlying systemic juvenile idiopathic arthritis. J Exp Med. 2017;214(11):3449–66.

33. Yang X, Garner LI, Zvyagin IV, Paley MA, Komech EA, Jude KM, et al. Autoimmunity-associated T cell receptors recognize HLA-B*27-bound peptides. Nature. 2022;612(7941):771–7.

34. Nelson MR, Tipney H, Painter JL, Shen J, Nicoletti P, Shen Y, et al. The support of human genetic evidence for approved drug indications. Nat Genet. 2015;47(8):856–60.

35. King EA, Davis JW, Degner JF. Are drug targets with genetic support twice as likely to be approved? Revised estimates of the impact of genetic support for drug mechanisms on the probability of drug approval. PLoS Genet. 2019;15(12):e1008489.

36. Ciccia F, Guggino G, Rizzo A, Alessandro R, Luchetti MM, Milling S, et al. Dysbiosis and zonulin upregulation alter gut epithelial and vascular barriers in patients with ankylosing spondylitis. Ann Rheum Dis. 2017;76(6):1123–32.

37. Guggino G, Mauro D, Rizzo A, Alessandro R, Raimondo S, Bergot AS, et al. Inflammasome activation in ankylosing spondylitis is associated with gut dysbiosis. Arthritis & Rheumatology. 2021;73(7):1189–99.

38. Ciccia F, Accardo-Palumbo A, Rizzo A, Guggino G, Raimondo S, Giardina A, et al. Evidence that autophagy, but not the unfolded protein response, regulates the expression of IL-23 in the gut of patients with ankylosing spondylitis and subclinical gut inflammation. Ann Rheum Dis. 2014;73(8):1566–74.

39. Yi K, Jo S, Song W, Lee HI, Kim HJ, Kang JH, et al. Single cell transcriptome and surface protein expression analysis identify OX40(+) GITR(+) pathogenic T helper 17 in ankylosing spondylitis. Arthritis Rheumatol. 2023.

