## Supplementary material for "Trained immunity in human monocyte enhances myeloid-T-cell pathogenic crosstalk in Ankylosing Spondylitis": Figure S1-S5

Supplementary tables

| Supplementary Table 1. Demographics of AS patients (n=51) | |
|---|---|
| Age, mean (range) years | 45 (25-89) |
| Sex, male/female | 32/19 |
| HLA-B27+ no. (%) | 34* (82.9%) |
| ESR, mean (range) | 11.8 (2-49) |
| CRP, mean (range) | 10.76 (0.2-87.5) |
| BASDAI, mean (range) | 4.3 (0.6-8.5) |
| NSAIDs therapy current | 28 (54.9%) |
| Biologic DMARD therapy current | 15 (29.4%) |
| Synthetic DMARD therapy current | 2 (4%) |
| * Unknown HLA-B27 status in 10 AS patients | |
| ESR: Erythrocyte sedimentation rate; CRP: C-reactive protein; BASDAI: Bath Ankylosing Spondylitis Disease Activity Index; NSAIDs: Non-steroidal anti-inflammatory drugs; DMARD: Disease-modifying antirheumatic drugs. | |

**Figure S1 scRNAseq analysis of unstimulated and LPS-stimulated PBMCs from three patients with AS.** (A) UMAP visualization of PBMCs colored by treatment conditions. (B) UMAP visualization of the major populations of all cells integrated from unstimulated and LPS-stimulated conditions. (C) Expression of a selected set of cluster markers. (D) UMAP visualization of monocytes colored by treatment conditions.

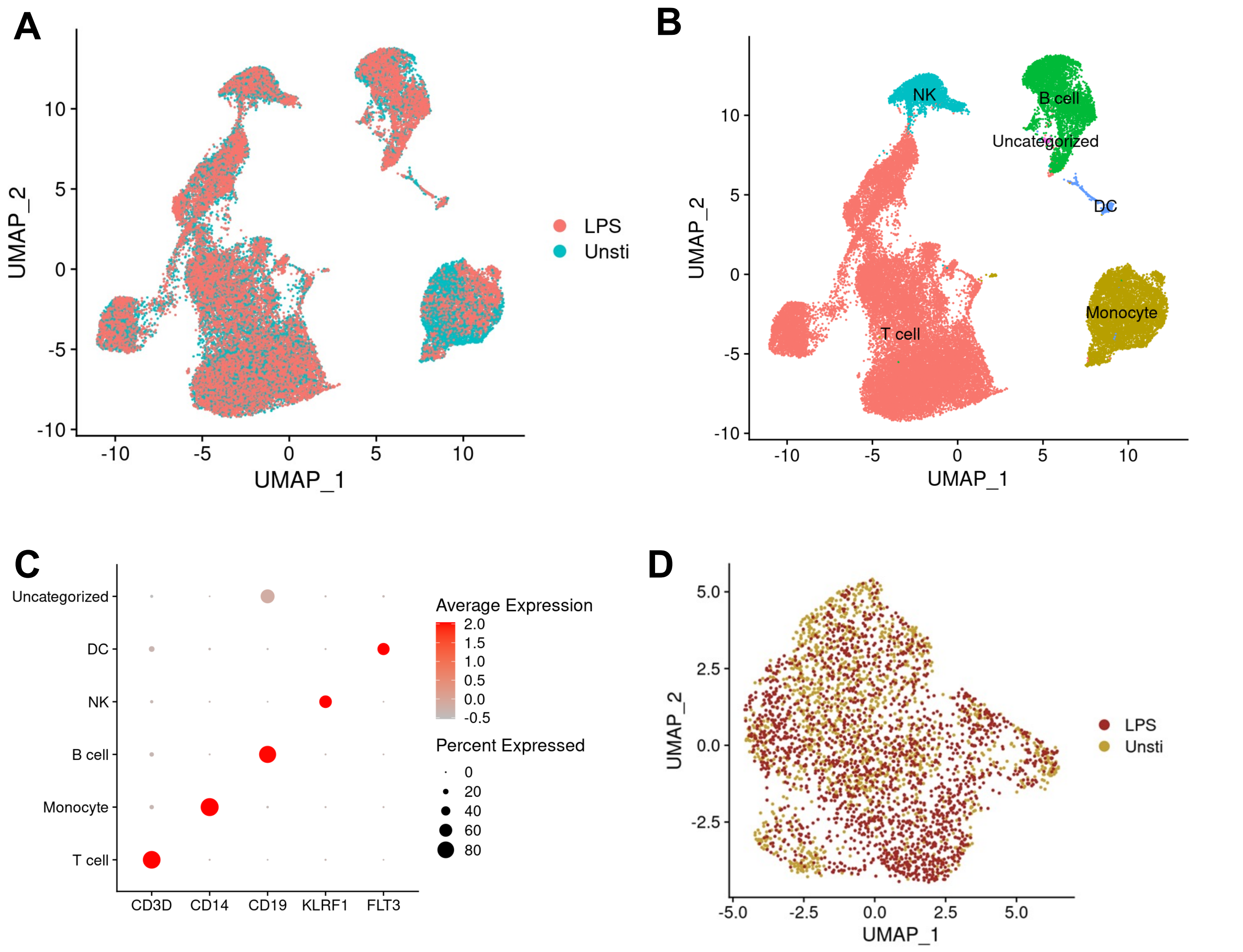

**Figure S2. T-cell activation beads (TABs) induce cytokine production by Ankylosing Spondylitis monocyte.** Gating strategy for identifying CD14+ monocytes in PBMCs of AS patients. Myeloid cells were gated on singlet cells and monocytes were defined as CD3−CD14+ cells.

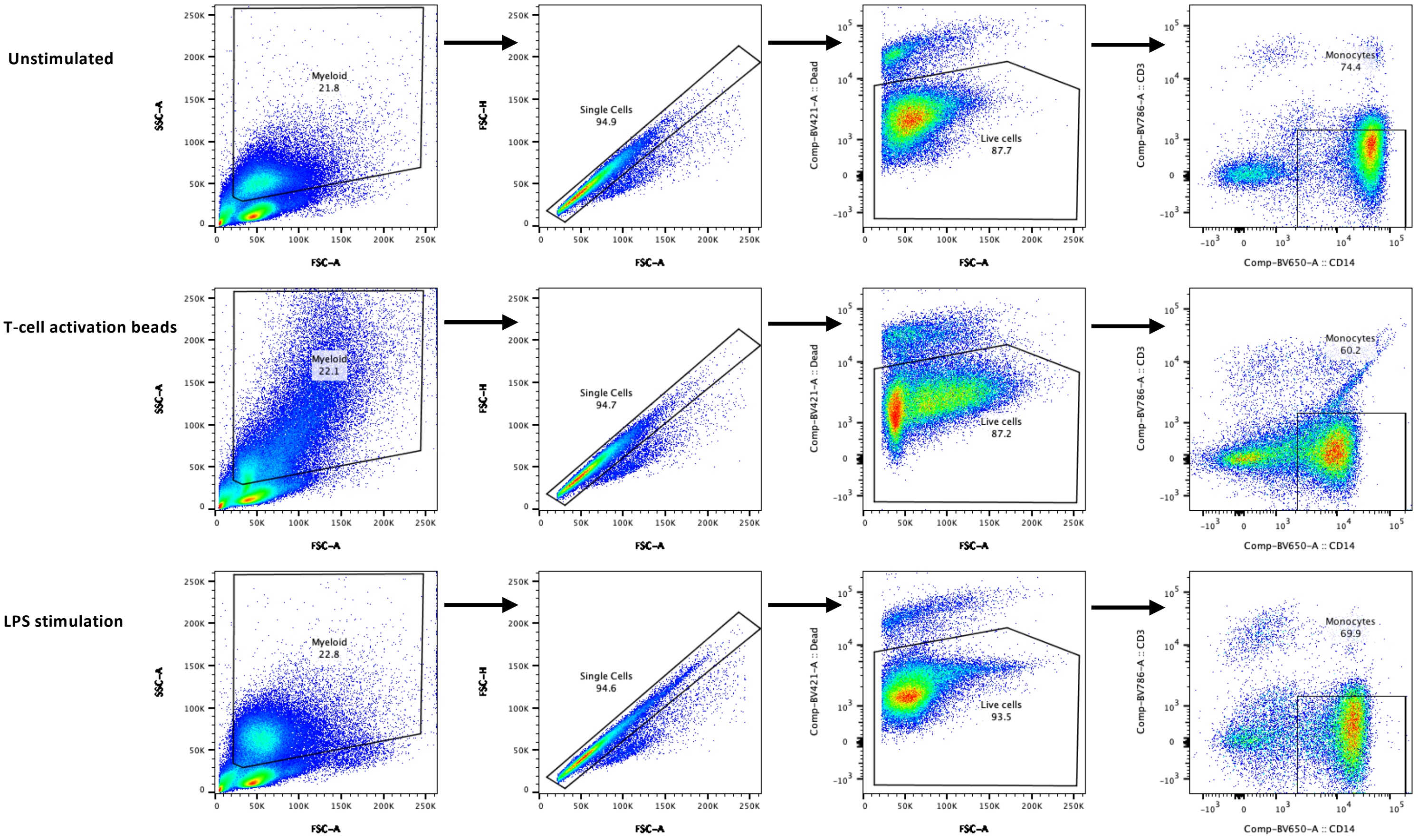

**Figure S3. scRNA sequencing analysis of T-cell activation bead (TAB)-stimulated PBMCs from three patients with AS.** (A) UMAP visualization of PBMCs from different conditions, colored by treatment conditions (TAB in orange, unstimulated in blue). Unstimulated dataset is same as in Fig S1. (B) UMAP visualization of the major populations of all cells integrated from unstimulated and TAB-stimulated conditions. (C) Expression of a selected set of cluster markers. (D) Expression of T cell activation genes induced by TAB stimulation

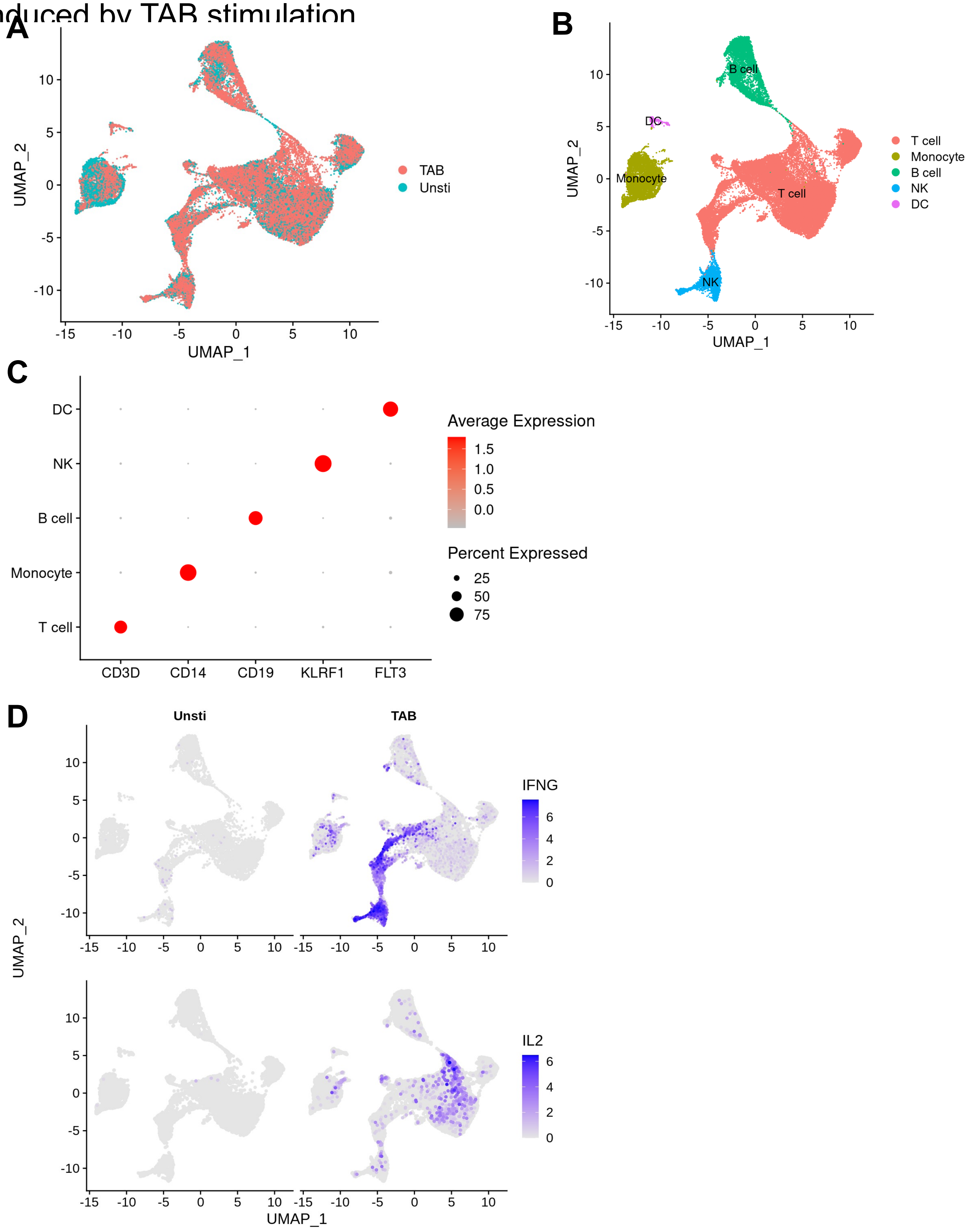

### Figure S4. Gating strategy for Th17 cells.

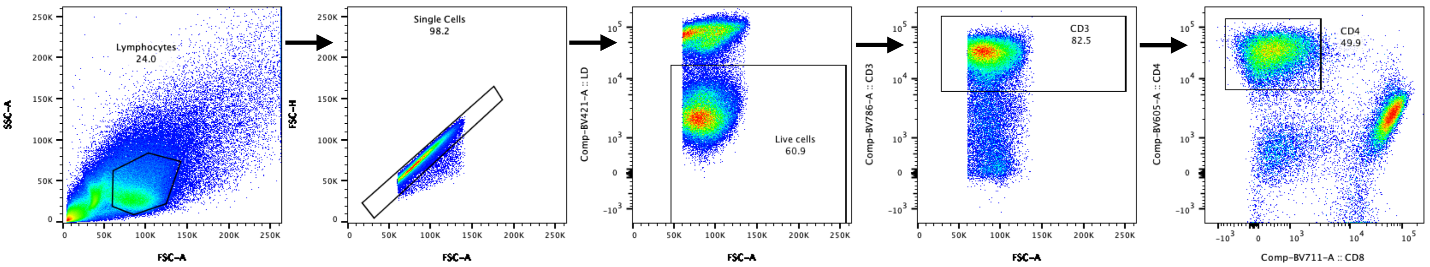

**Figure S5. Annotation of T-cell subsets from matched blood and synovial fluid from patients with AS**. Expression of marker genes in different clusters.

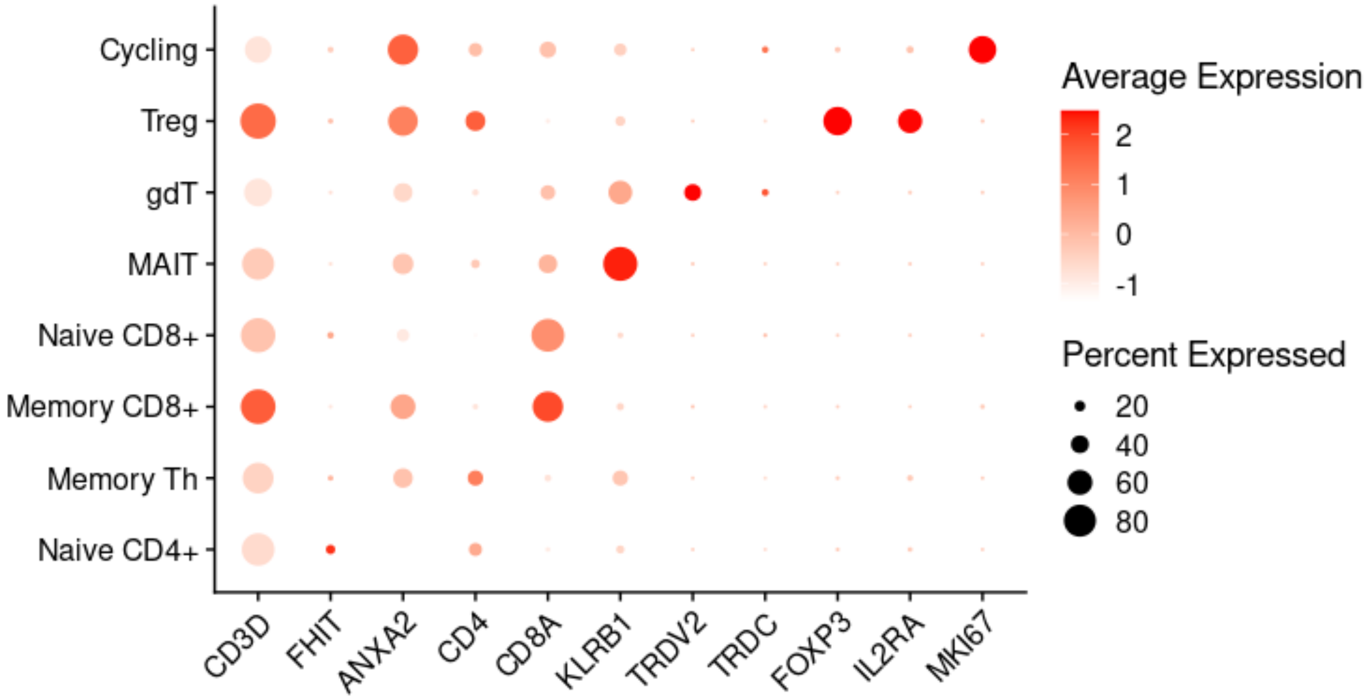
